# Genoppi: a web application for interactive integration of experimental proteomics results with genetic datasets

**DOI:** 10.1101/115576

**Authors:** April Kim, Edyta Malolepsza, Justin Lim, Kasper Lage

**Affiliations:** Broad Institute of MIT and Harvard, Cambridge, MA 02142, USA; Department of Surgery, Massachusetts General Hospital, Boston, MA 02142, USA; Massachusetts Institute of Technology, Cambridge, MA 02142, USA

## Abstract

**Summary:** Integrating protein-protein interaction experiments and genetic datasets can lead to new insight into the cellular processes implicated in diseases, but this integration is technically challenging. Here, we present Genoppi, a web application that integrates quantitative interaction proteomics data and results from genome-wide association studies or exome sequencing projects, to highlight biological relationships that might otherwise be difficult to discern. Written in R, Python and Bash script, Genoppi is a user-friendly framework easily deployed across Mac OS and Linux distributions.

**Availability:** Genoppi is open source and available at https://github.com/lagelab/Genoppi

## 1 Introduction

With recent advances in stem cell technologies and quantitative proteomics methods, it is now possible to experimentally interrogate the physical interactions of proteins in a tissue-or cell-type-specific manner at scale. In parallel, the ongoing genomic revolution has enabled the identification of common variant loci significantly associated with diseases through genome-wide association studies (GWAS). Exome sequencing technologies have also led to the identification of specific genes or protein-coding mutations linked to particular diseases. Since it has been shown that genes implicated in a common complex traits often interact at the level of proteins (Rossin et al., 2011; Lundby et al., 2014; Lage, 2014) it is desirable to integrate the results of quantitative proteomics experiments and user-defined genetic datasets in an interactive workflow that can be shared with collaborators.

Towards that aim, we developed an open-source application Genoppi. It provides (i) quality control (QC) of mass spectrometry-based interaction proteomics datasets and (ii) on-the-fly integrative analyses of the proteomic data and user-defined GWAS or exome sequencing studies. By allowing users to store the results of analyses and visualizations, Genoppi facilitates data sharing in cross-disciplinary collaborations.

## 2 Features

In Genoppi, users can upload experimental quantitative interaction proteomics data from immunoprecipitations (IP) or full proteome analysis followed by tandem mass spectrometry (MS/MS). Upon loading a file, the software plots interactive graphical representations of the data as volcano and scatter plots (Fig. 1). As part of the QC workflow, known human protein interaction partners from InWeb (Rossin et al., 2011; Lage, 2014; Li et al., 2016) can be identified. InWeb contains known protein-protein interactions from >40,000 articles and overlaying this information with the experimental data enables users to easily distinguish new interactions from those already reported in the literature.

**Fig. 1.**
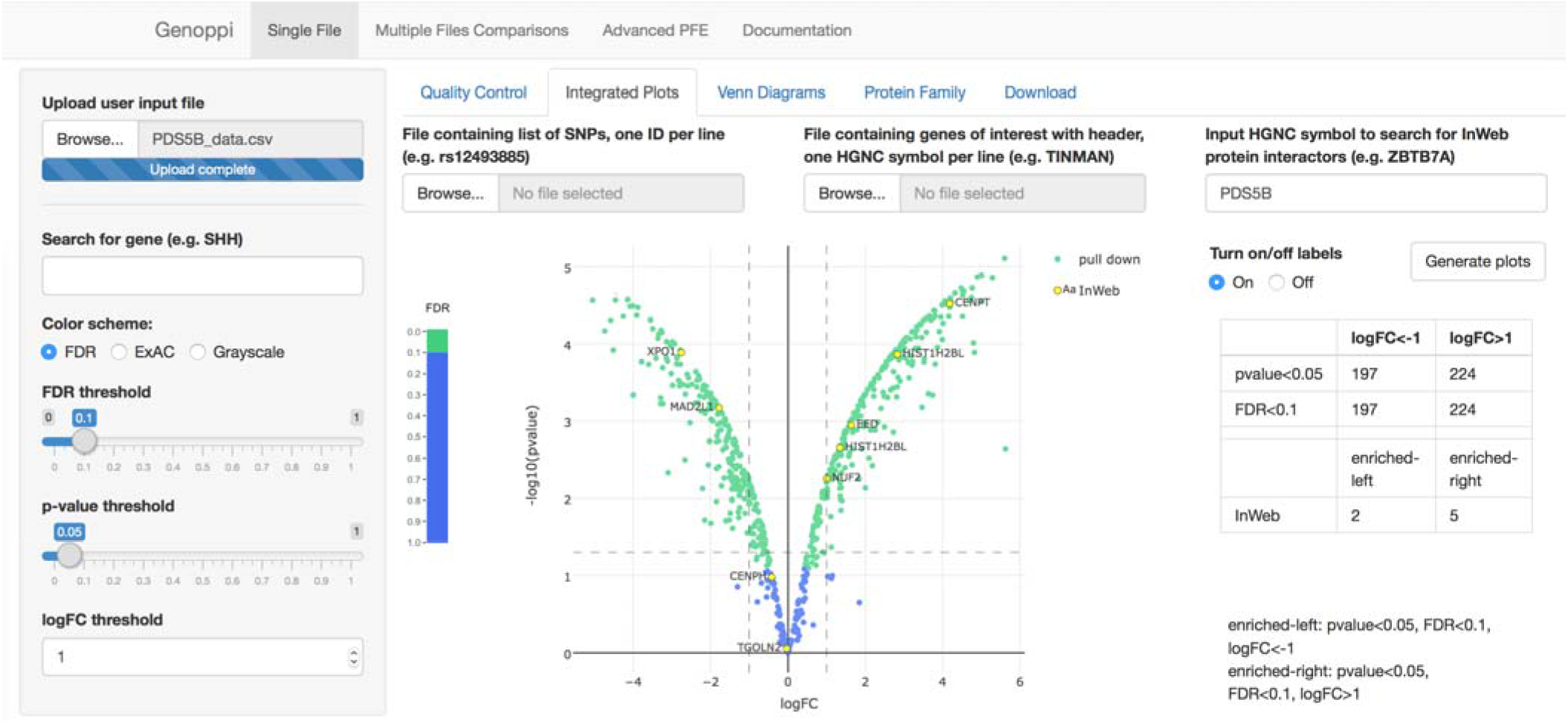
Genoppi interface. Figure shows a volcano plot of mock proteomics results. Log_2_ fold change (logFC) values of identified proteins are plotted on the x-axis and the negative log_10_ transformed P values of that enrichment on the y-axis. Proteins with a false discovery rate (FDR) < 0.1 are green, and those with FDR >= 0.1 are blue. Known interaction partners of the bait determined by InWeb are highlighted in yellow. Different color schemes are available: grayscale and the intolerance to loss-of-function variation (pLI) of a given gene.

Further, Genoppi allows users to (i) overlay GWAS data by inputting single nucleotide polymorphisms (SNP), (ii) map UniProt identifiers (The UniProt Consortium, 2015) to HUGO Gene Nomenclature Committee (HGNC) symbols (Gray et al., 2015), (iii) execute protein family (PFE) analyses, (iv) identify proteins encoded by genes intolerant to loss-of-function mutations as determined by the Exome Aggregation Consortium (ExAC) data (Lek et al., 2016), and (v) highlight specific proteins corresponding to users’ uploaded genes of interest (e.g., a set of genes from the Genotype Tissue Expression [GTEx] project).

When different datasets are integrated with proteomics results, overlaps are statistically tested and visualized. For example, the overlap between interaction partners of bait identified in MS/MS experiment and genes linked to a particular disease is determined.

A significant barrier to using the results from GWAS is that SNP-to-gene mapping is nontrivial. For this reason, the application automatically maps SNPs to genes using haplotype information from 1KGP (The 1000 Genomes Project Consortium, 2015) and highlights those genes in the proteomic data. If multiple protein-coding genes are present in genetic locus defined by a SNP of interest, all genes are mapped as candidate genes for the associated SNP. Together, this allows users to identify interaction partners of bait that are encoded in loci genetically associated with phenotype relevant to the bait.

PFE analyses can be applied to either single bait versus control experiment or comparison between different experimental conditions (e.g., with and without drug treatment on cells with various mutations). The latter format allows users to identify human protein families defined by proteins enriched in the conditions of choice that can serve as an additional QC or to guide hypothesis generation or follow up experiments.

Users can download generated data in text format and interactive plots as HTML reports. Genoppi can be archived and easily modified to suit users’ custom needs.

## 3 Implementation

Genoppi is written using R standard packages (R Core Team, 2016) using Shiny framework (Chang et al., 2016) and Bash script. Automated assignment of human UniProt accession numbers to HGNC symbols is written in Python. Median normalization of user dataset and single sample moderated t-test is performed using Bioconductor limma package (Ritchie et al., 2015). Pairwise measure of LD was calculated using VCFtools (Danecek et al., 2011) on VCF files containing genotype data from phase 3 of 1KGP. R graphing library plotly (Sievert et al., 2016) was used to create interactive graphs.

## Acknowledgements

EM thanks Zuzana Tothova, Josephine Kahn, Siddhartha Jaiswal, and Srinivas Viswanathan for discussion on data analysis, and the Proteomics Platform at the Broad Institute for explaining output structures.

## Author Contribution

AK, EM, JL developed algorithms with supervision from KL. AK developed Genoppi application. AK, EM, KL wrote paper. KL initiated and led project.

